# Experimental support for human papillomavirus genome amplification early after infectious delivery

**DOI:** 10.1101/2022.06.24.497572

**Authors:** Katarzyna Zwolinska, Malgorzata Bienkowska-Haba, Rona S. Scott, Martin Sapp

## Abstract

Even though replication and transcription of human papillomavirus type 16 (HPV16) has been intensively studied, little is known about immediate early events of the viral life cycle due to the lack of an efficient infection model allowing genetic dissection of viral factors. We employed the recently developed infection model (Bienkowska-Haba et al. PLOS Pathogen 2018) to investigate genome amplification and transcription immediately after infectious delivery of viral genome to nuclei of primary keratinocytes. Using EdU pulse labeling and highly sensitive fluorescence *in situ* hybridization, we observed that the HPV16 genome is replicated and amplified in an E1 and E2 dependent manner. Knockout of E1 resulted in failure of the viral genome to replicate and amplify. In contrast, knockout of the E8^E2 repressor led to increased viral genome copy number confirming previous reports. Genome copy control by E8^E2 was confirmed for differentiation-induced genome amplification. Lack of functional E1 had no effect on transcription from the early promoter, suggesting that viral genome replication is not required for p97 promoter activity. However, infection with an HPV16 mutant virus defective for E2 transcriptional function revealed a requirement of E2 for efficient transcription from the early promoter. In absence of the E8^E2 protein, early transcript levels are unaltered and even decreased when normalized to genome copy number. Surprisingly, lack of functional E8^E2 repressor did not affect E8^E2 transcript levels when normalized to genome copy number. These data suggest that the main function of E8^E2 in the viral life cycle is to control genome copy number.

**IMPORTANCE:** It is being assumed that HPV utilizes three different modes of replication during its lifecycle: initial amplification during the establishment phase, genome maintenance and differentiation-induced amplification. However, HPV16 initial amplification was never formally proven due to a lack of an infection model. Using our recently established infection model (Bienkowska-Haba et al. PLOS Pathogen 2018), we demonstrate herein that viral genome is indeed amplified in an E1 and E2 dependent manner. Furthermore, we find that the main function of the viral repressor E8^E2 is to control viral genome copy number. We did not find evidence that it regulates its own promoter in a negative feedback loop. Our data also suggest that the E2 transactivator function is required for stimulation of early promoter activity, which has been debated in the literature. Overall, this report confirms the usefulness of the infection model for studying early events of the HPV lifecycle using mutational approaches.

## INTRODUCTION

Human papillomaviruses (HPV) are small non-enveloped dsDNA viruses infecting basal keratinocytes of the stratified cutaneous or mucosal epithelia causing benign and malignant outgrowths (1–3). HPV establish infection by delivery of its chromatinized, circular double-stranded DNA genome to the nucleus of basal keratinocytes, where the viral genome is residing in an episomal form (2). The current assumption is that viral genome is amplified after infectious delivery to the nucleus of target cells to establish infection. After which the genome copy number is maintained. When HPV harboring cells enter the terminal differentiation stage, viral genome is massively amplified prior to capsid protein synthesis and progeny virion production (4). Differentiation-induced viral genome amplification to high copy number requires recombination-dependent replication (RDR) supported by the DNA damage response (DDR) (2, 4–6). These sequences of events suggest three modes of replication are required to complete the HPV life cycle.

Two viral factors are essential for viral genome replication. The E1 protein is essential for initiating genome replication and for recruitment of the host cell replication machinery. In addition to its function in replication initiation, it unwinds the double-stranded DNA during elongation. It is subject to various post-translational modifications that are thought to regulate its activity (7). The E2 protein forms a complex with E1 protein to direct its binding to the origin of replication. E2 protein is also a transcriptional regulator, which acts by recruiting inhibitory or stimulatory cellular factors to the viral genome (8). E2 protein also tethers viral genomes to mitotic chromosomes during mitosis through interactions between viral DNA and the carboxyl terminal E2 DNA binding domain on one side, and the E2 transactivation domain and specific proteins in the mitotic chromosomes to ensure proper partitioning of viral genome to daughter cells. One of the best-known targets for E2 is the chromatin adapter protein, Brd4, which has been implicated in viral genome tethering for some but not all HPV types (9, 10). The replication and transcriptional regulation function of E2 can be separated by mutations in the amino terminal transactivating domain. I73A or R37A substitutions cause abrogation of the transcriptional activity of E2 but leave its replication functions intact. However, Brd4 binding is impaired and results in loss of viral genome over time because of incorrect tethering and uneven partitioning (8, 11). The E39A mutation in HPV16 E2 abrogates E1 binding and results in a viral replication defect, while preserving the transactivation functions (8, 12, 13). Introduction of mutations in conserved regions of the E2 DNA binding domain interferes with E1-E2 binding to the viral origin of replication and to enhancer and promoter elements located in the long control region thus impairing both replication and transcription functions (14–16).

Another viral protein, E8^E2, has been shown to regulate genome copy number. This protein arises from a splice product of the E8 gene located in E1 open reading frame spliced to the carboxyl terminus of E2 comprising the hinge and DNA binding domain of the E2 gene. Expression of E8^E2 is driven by the pE8 promoter, which is in the E1 ORF approximately 70-150 nt upstream of the E8 start codon (17). Knock out of HPV16 E8^E2 resulted in 10-100 times higher viral genome levels in normal or immortalized keratinocytes (17) as well as in organotypic raft cultures of stable keratinocytes cell lines (18). HPV16 E8^E2 knock out genomes were maintained long-term as extrachromosomal elements, whereas HPV31 ΔE8^E2 DNA integrated into host genomes (17, 19, 20). E8^E2 is also known as repressor of early and late promoter expression (18, 19), as well as a negative regulator of its own promoter (17, 21). The mechanism of repression of E1/E2 dependent viral DNA replication and viral transcription may be due to their competing for the same binding site and the formation of inactive E2/E8^E2 heterodimers (19, 22). It was also shown that the E8 domain interacts with NCOR1/SMRT repressor complexes consisting of GPS2, HDAC3, and TBl1 and TBLR1 proteins (17, 22, 23).

Some inconsistencies exist regarding the exact role of E1 and E2 protein especially during the maintenance stage. Some reports suggest an E1-independent viral genome synthesis during genome maintenance in undifferentiated keratinocytes (24, 25). Simultaneous E1-dependent and E1-independent amplification during the maintenance and vegetative stage of infection has also been reported (6). Knock out or knock down of E1 protein prevents viral DNA amplification (6, 24). These inconsistencies warrant additional studies.

Even though it has been generally assumed that viral genome is amplified after infectious entry into keratinocytes, this assumption has never been experimentally supported and was only derived from copy number measurements in established cell lines. The lack of experimental support was largely due to a lack of good infection models. Most of the present knowledge about papillomaviruses replication is based on models using transfection of human epithelial cell lines (6, 15, 26–30) or primary human keratinocytes (31, 32). We now took advantage of our recently established infection model using an ECM-to-cell transfer approach to investigate the fate of viral genome after infectious entry (33). Furthermore, there are conflicting reports regarding the roles of E1 and E2 during the establishment and maintenance stages of the HPV life cycle. The role of the E8^E2 protein during the full viral life cycle was never confirmed in the context of infection. In the present study, we demonstrate immediate E1 and E2-dependent viral genome amplification in infected human primary foreskin keratinocytes (HFK) and confirm the involvement of the E8^E2 protein as regulator of viral genome copy number. However, E8^E2 knockout did not affect viral transcript levels whereas the E2 transactivation function was required for efficient transcription immediately after infectious delivery.

## RESULTS

### Initial amplification of HPV16 genome requires presence of both E1 and E2 proteins

To evaluate the role of E1 and E2 proteins in the initial amplification of viral genome, we infected primary human foreskin keratinocytes (HFK) through ECM-to-cell transfer with HPV16 WT as well as E1-TTL, E2-E39A/R302G, and E2-I73A mutated quasivirions as previously described (33). A similar multiplicity of infection (MOI, 100-200 vge/cell) was used for all viruses based on qPCR measurement of encapsidated viral genome. Both, E1-TTL and E2-E39A/R302G mutant viruses are defective for viral replicative functions. While the E1-TTL mutant has a translation termination linker inserted at position 892-894, the E2-E39A/R302G carries two separate single point mutations (2871A/C and 3659A/G). The mutant protein retains Brd4 interaction functions but is defective for supporting viral replication as well as transactivation (8). The E2-I73A mutant has substituted two nucleotides in the transactivation region (2972-73AT/GC) resulting in impaired transcriptional regulation and Brd4 binding. Infected cells as well as mock-infected control cells were cultured in the presence of Rho kinase (ROCK) inhibitor Y-27632 and grown in the presence of mouse fibroblasts as feeder cells for HFK. At 1-, 2-, 4- and 7-days post infection (dpi), keratinocytes were collected, and total DNA was isolated. Viral genome was measured by qPCR with primers specific for HPV16 E6 and normalized to the cellular housekeeping gene cyclophilin A (CyPA) (Fig 1A-B). There was no significant difference in HPV16 genome level at 2 dpi (Fig 1A) in WT, E1-TTL and E2-E39A/R302G infected keratinocytes. This early post infection, viral DNA from the inoculate masks any replication differences, however, data confirm that keratinocytes were infected at a similar MOI. Genome copy numbers are significantly higher in established HPV16-immortalized keratinocytes. Continued culturing of infected cells revealed increasing levels of viral DNA in WT infected cells and significant losses in both E1-TTL and E2-E39A/R302G infected HFK.

**FIG 1.**
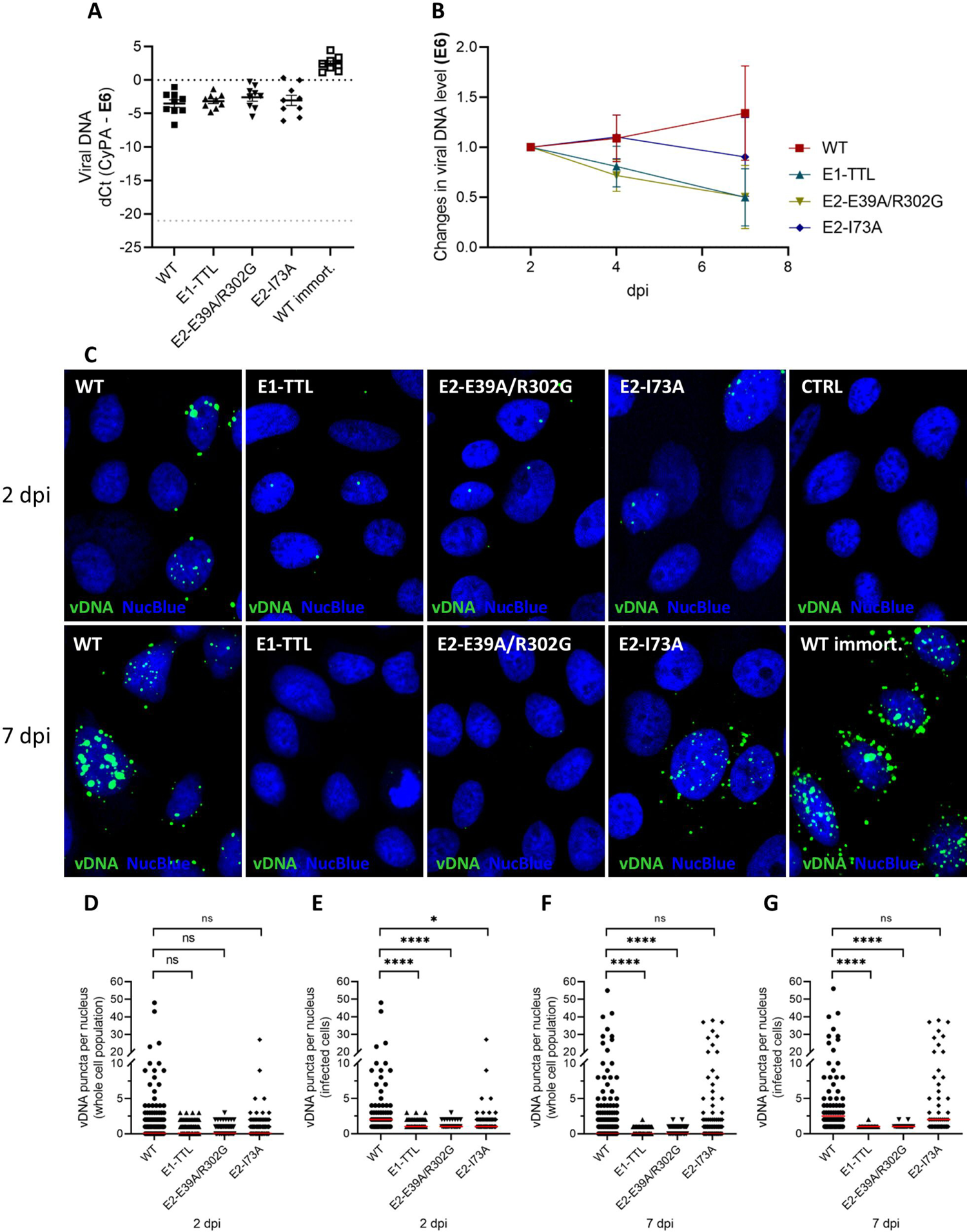
Amplification of viral genome early after infection of human foreskin keratinocytes (HFKs) with HPV-16 WT, E1-TTL, E2-E39A/R302G, and E2-I73A mutants, in monolayer culture. (A) Viral genome level as measured using E6 specific primers at 2 dpi shown as dCt normalized to the cyclophilin A (CyPA). Viral DNA in WT-immortalized cells shown for comparison. Light gray dot line shows limit of detection. (B) Changes in viral genome (E6) level over 7 dpi. The data shown on graph are fold changes relative to results for 2 dpi, normalized to CyPA. Error bars represent SEM of 9 independent experiments. (C) Detection of viral DNA in infected HFKs growing in monolayer culture, 2 and 7 dpi, by highly sensitive fluorescent hybridization. (D-G) Quantification of viral DNA puncta detected by DNAscope in nuclei of infected keratinocytes 2 and 7 dpi. (D and F) Approximately 200 cells in each group (between 196-224) were used for the analysis of nuclear viral DNA signal. All cells, positive and negative for viral genome puncta were analyzed. (E and G) Only cells with nuclear DNA scope signal were included in the analysis. Data presented as individual values with median (red line). Statistically significant differences are marked by asterix (*p<0.05 and ****p<0.0001).

The increase and decrease, respectively, was consistently observed in all but one biological replicates. The high variability resulting in large error bars is due to cell growth differences of primary keratinocytes due to donor variation and infection efficiency. Viral DNA level decreased slower in cells infected with E2-I73A mutant viruses (Fig 1B). Viral genomes in WT and mutant infected cells were also visualized using DNA hybridization with HPV16/18 L1-specific probes (“DNAscope”). We chose an L1 specific set of probes for these analyses since late transcripts are at the limit of detection in our undifferentiated cultures. The methodology employed allows the detection of single viral genomes without detecting viral transcripts (Fig. 2). Uninfected cells were used as negative and HPV16 immortalized cells as positive controls (Fig. 1C). At 2 dpi, we observed viral genomes delivered to nuclei of approximately 20% of cells infected with WT, E1-TTL, E2-E39A/R302G, and E2-I73A. Amplification foci were observed in some WT-infected cells as well as some I73A at 2 dpi and were much more pronounced at 7 dpi (Fig. 1C). Larger genome foci suggested that viral genome is amplified immediately after infectious delivery to the nucleus. Only single genome copies were detected in cells infected with E1-TTL and E2-E39A/R302G mutant viruses at 2 dpi. At 7 dpi, only few cells retained HPV16 E1-TTL and E2-E39A/R302G mutant genome. Quantification of DNA scope data confirmed a statistically significant difference between WT and E1-TTL and E2-E39A/R302G mutant viruses at 2 and 7 dpi (Fig. 1D-G). Amplification foci were also seen in some E2-I73A-infected cells 2 as well as 7 dpi. While the number of genome foci was not statistically different, the number of genome harboring cells was lower in E2-I73A infected cells suggesting a defect in genome maintenance. We consistently observed cytosolic localization of some viral genome in WT and E2-I73A infected cells at 7 dpi. Controls suggest that cytosolic signals are not due to detection of viral mRNA or unspecific staining of host cell RNAs (see Fig. 2 for controls).

**FIG 2.**
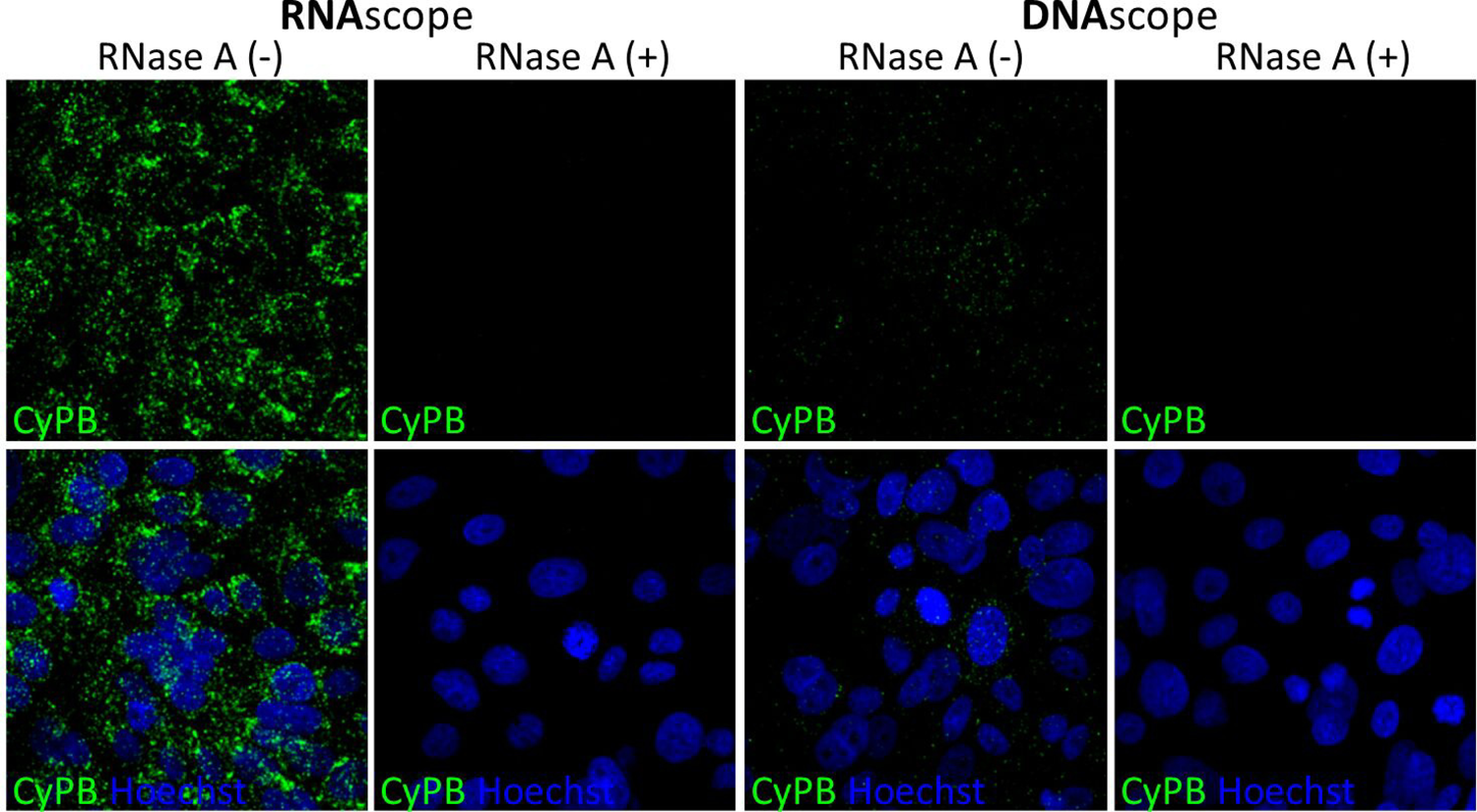
Controls for specificity of highly sensitive *in situ* hybridization of RNA (RNAscope) and DNA (DNAscope). Uninfected human foreskin keratinocytes were subjected to hybridization with the probe for human cyclophilin B (CyPB) in RNAscope (left panels) and DNAscope (right panels) conditions, in the presence or absence of RNase A.

To confirm that viral HPV16 genome is indeed replicated and amplified immediately after infectious delivery, we employed EdU labeling. At 12 to 16 hours post infection, 10µM of 5-ethynyl-2’-deoxyuridine (EdU) was added to the media to label newly synthesized DNA. EdU was omitted from controls. At 24 or 48 hpi, cells were harvested, lysed, and isolated DNA was subjected to biotinylation with PEG biotin azide using Click-iT chemistry. Biotinylated DNA was captured by streptavidin-coated magnetic beads. Total DNA and EdU-DNA was quantified using qPCR with HPV16 E6 and cellular cyclophilin A (CyPA) specific primers. EdU labeled DNA level was expressed as difference between Ct values of DNA pulled down from unlabeled and labeled samples (dCt) normalized to input DNA. The level of infection under the different assay conditions was comparable as measured by HPV16 E7 mRNA using RT-qPCR (Fig 3A-B). At 1 and 2 dpi, we were able to detect EdU-labeled cellular DNA (CyPA) in all infected and non-infected control cultures. EdU-labeled viral DNA was detected only after infection with WT HPV16 and E2-I73A, but not with E1-TTL and E2-E39A/R302G mutant viruses (Fig 3C-D). At one dpi, the level of newly synthesized viral DNA was similar in WT and E2-I73A but dropped to ∼75% in E2-I73A mutant virus infected cells at 2 dpi. Since we are employing an infection model for nuclear delivery of viral genome, only a fraction of cells was infected at 1 and 2 dpi (approximately 5 to 30% of keratinocytes). This likely explains why the pull down of the host cell control CyPA shows higher differentials than the viral DNA. It does not imply that viral genome is less efficiently replicated than the host gene. Taken together, these results strongly suggest that viral genome is replicated and amplified after infectious delivery, in an E1 and E2 protein dependent manner.

**FIG 3.**
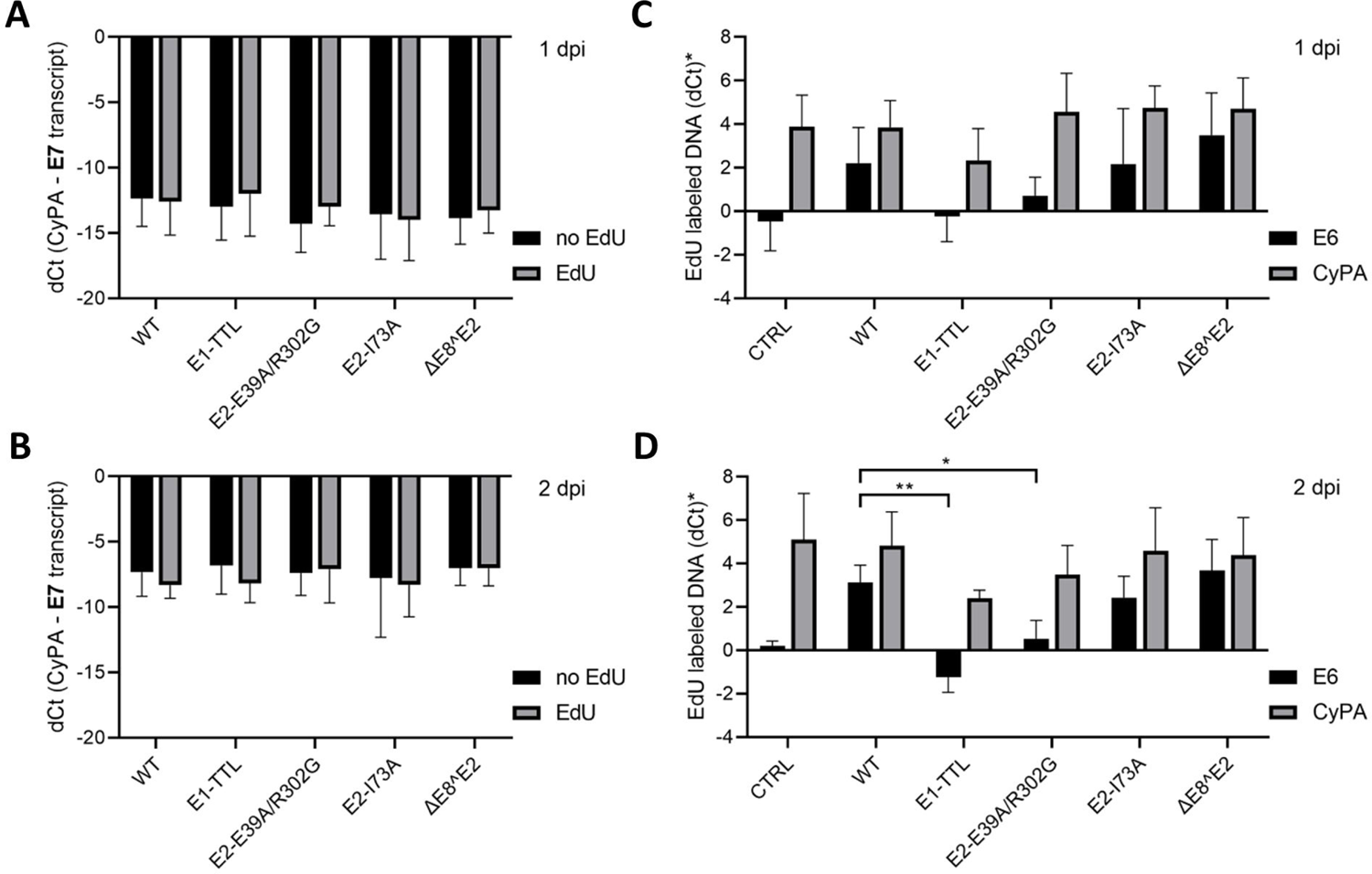
Detection of EdU-labeled amplifying viral genome in HPV-16 WT, E1-TTL, E2-E39A/R302G, E2-I73A, and ΔE8^E2 mutant infected HFKs immediate early after infection. (A, B) Viral transcripts (E7) in EdU-treated and untreated HFKs at 1 and 2 dpi, respectively. Transcripts expressed as dCt relative to the cyclophilin A (CyPA). Error bars represent SEM of 3 (1 dpi) or 5 (2 dpi) independent experiments. (C, D) Viral (E6) and cellular (CyPA) EdU labeled DNA at 1 and 2 dpi respectively. dCt* = unlabeled DNA dCt (pull down – input) – EdU labeled DNA dCt (pull down – input) Differences statistically significant at *p<0.05 and **p<0.01.

### Genome copy number regulation by E8^E2 protein during establishment of HPV16 infection

To assess involvement of E8^E2 protein in early viral amplification, we generated ΔE8^E2 HPV16 quasiviruses carrying a single nucleotide substitution 1281G/A in the E8 open reading frame. The mutation results in disruption of E8 KWK motif by introducing a stop codon resulting in the functional knock out of the E8^E2 protein (18). We infected primary human keratinocytes with WT and ΔE8^E2 mutated quasiviruses through ECM-to cell transfer (33). Average viral DNA level, measured by qPCR, was around 25% higher at 1 dpi in ΔE8^E2 than WT infected cells and this difference increased over time. Starting at 2 dpi the difference in viral genome levels between WT and ΔE8^E2 was statistically significant (Fig 4A, B) and was 2.5 times higher in ΔE8^E2 at 7 dpi. To verify that the observed difference was a result of viral amplification, we performed EdU labeling and pull down of newly synthesized DNA, at 1 and 2 dpi as described above (Fig 3C, D, respectively). Nascent viral DNA level was between 18-58% higher in ΔE8^E2 compared to WT infected cells at both, one and two dpi. We also visualized HPV16 genome amplification in ΔE8^E2 and WT infected cells at 1, 2 and 7 dpi using DNAscope (Fig 4C). We were able to detect incoming WT and ΔE8^E2 viral genome puncta delivered to nuclei of infected keratinocytes and larger replication foci in nuclei of some cells at 1 dpi. Viral DNA foci were more numerous and larger in ΔE8^E2 infected cells already at 2 dpi (Fig. 4D-G). Again, we find that some viral genome localizes to the cytoplasm. Cytoplasmic localization is more pronounced in cells also containing large quantities of nuclear genome. Taken together, these data support an E8^E2 role in regulation of E1/E2 dependent genome amplification early after infection.

**FIG 4.**
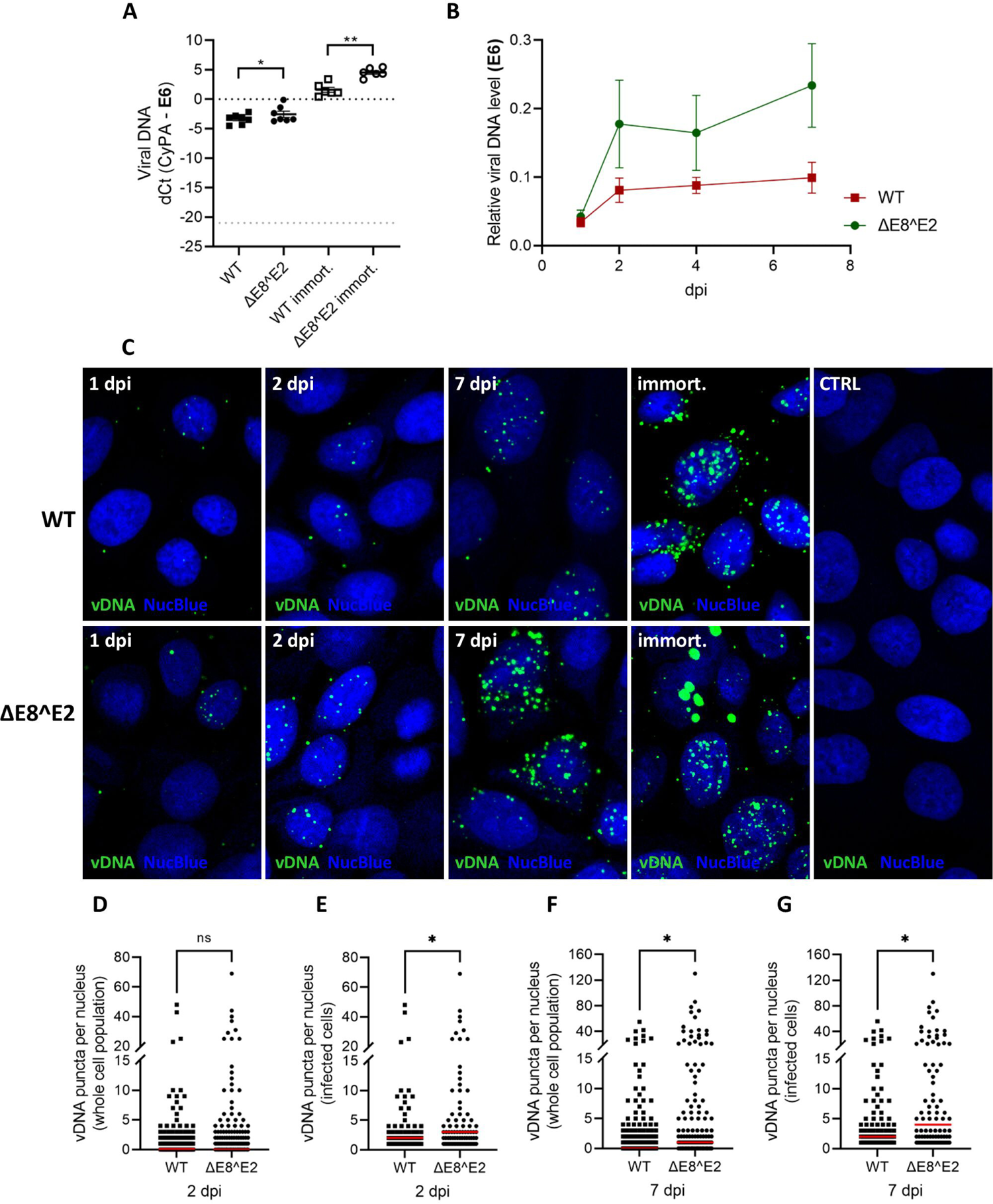
Amplification of viral genome early after infection of human foreskin keratinocytes (HFKs) with HPV-16 WT and ΔE8^E2 mutants, in monolayer culture. (A) DNA (E6) level at 2 dpi shown as dCt normalized to the cyclophilin A (CyPA). Viral DNA in WT-, and ΔE8^E2-immortalized cells shown for comparison. Light gray dot line shows limit of detection. (B) Viral genome (E6) relative level normalized to CyPA over 7 dpi. Error bars represent SEM of 7 independent experiments. (C) Detection of viral DNA in infected HFKs growing in monolayer culture, 1, 2, 7 dpi and in immortalized cells, by highly sensitive fluorescent hybridization. (D-G) Quantification of viral DNA puncta detected by DNAscope in nuclei of infected keratinocytes 2 and 7 dpi. (D and F) Approximately 200 cells each (between 200-224) were used for the analysis independent of nuclear DNA scope signal. (E and G) Only cells with nuclear DNA scope signal were included in the analysis. Data presented as individual values with median (red line). Statistically significant differences are marked by asterix (*p<0.05 and ****p<0.0001).

### Immediate/early viral transcription does not require viral genome amplification

We also took advantage of the mutant viruses to explore the contribution of genome replication to transcriptional regulation after infectious delivery of viral genome. To this end, primary human foreskin keratinocytes were infected through ECM-to-cell transfer with HPV16 WT as well as E1-TTL, E2-E39A/R302G, E2-I73A, and ΔE8^E2 quasivirions. At 1, 2, 4 and 7 dpi, keratinocytes were harvested for measuring viral transcripts using RT-qPCR. Viral transcript levels were normalized to the cellular housekeeping gene, cyclophilin A (CyPA). Expression of viral RNA was also visualized by using RNA hybridization (RNAscope) with a probe targeting HPV16/18 E6/E7. We did not observe significant differences in early (E6, E7, E1, E2) and E1^E4 transcript levels in keratinocytes infected with WT and mutants at 2 dpi (Fig 5A-E; 6A-F). However, carrying infection on for 4 and 7 days revealed a drastic drop of viral transcripts in both E1-TTL and E2-E39A/R302G infected cells compared to WT infected cells. The reduction of early transcript levels was not as pronounced and occurred slower in E2-I73A mutant infected cells (Fig 5F). This trend was also observed in RNA scope results (Fig 5G). We conclude that initial viral transcription does not require genome replication. Subsequent decrease of transcript levels at later times post infection in E1-TTL and both E2 mutant virus infected cells is likely due to a loss of viral genome. As expected, early viral transcripts increased over time in ΔE8^E2 infected cells similar to WT up to 7 dpi (Fig 6G-H). Normalization of mRNA to the genome level did not show significant differences in ΔE8^E2 and WT infected cells during the first 7 dpi (Fig 7D1-7D3). However, a consistently lower level of p97 derived transcripts in E8^E2 mutant infected cells was observed. These data suggest that E8^E2 may not repress the early p97 promoter early in infection as previously indicated.

**FIG 5.**
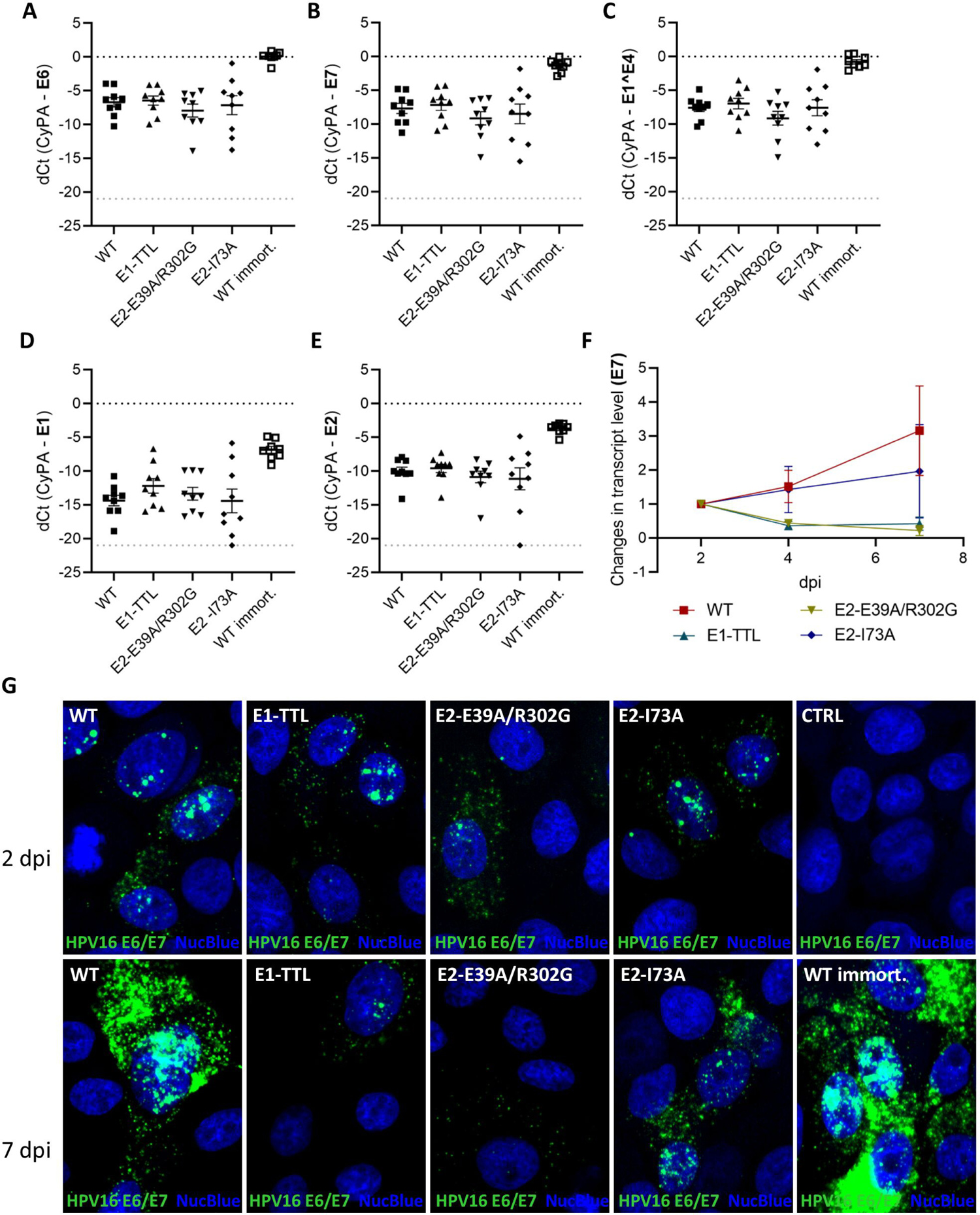
Expression of viral transcripts in HPV16 WT, E1-TTL, E2-E39A/R302G, and E2-I73A mutant infected HFK early after infection. E6, E7, E1^E4, E1, E2 levels at 2 dpi are shown as dCt relative to the housekeeping control gene cyclophilin A (CyPA) (A-E respectively). Viral transcripts in WT-immortalized cells shown for comparison. Light gray dot line indicates limit of detection. (F) Changes in E7 transcript relative to results for 2 dpi, normalized to CyPA. Error bars represent SEM of 9 independent experiments. (G) Representative RNAscope images of infected HFKs growing in monolayer culture at 2 and 7 dpi.

**FIG 6.**
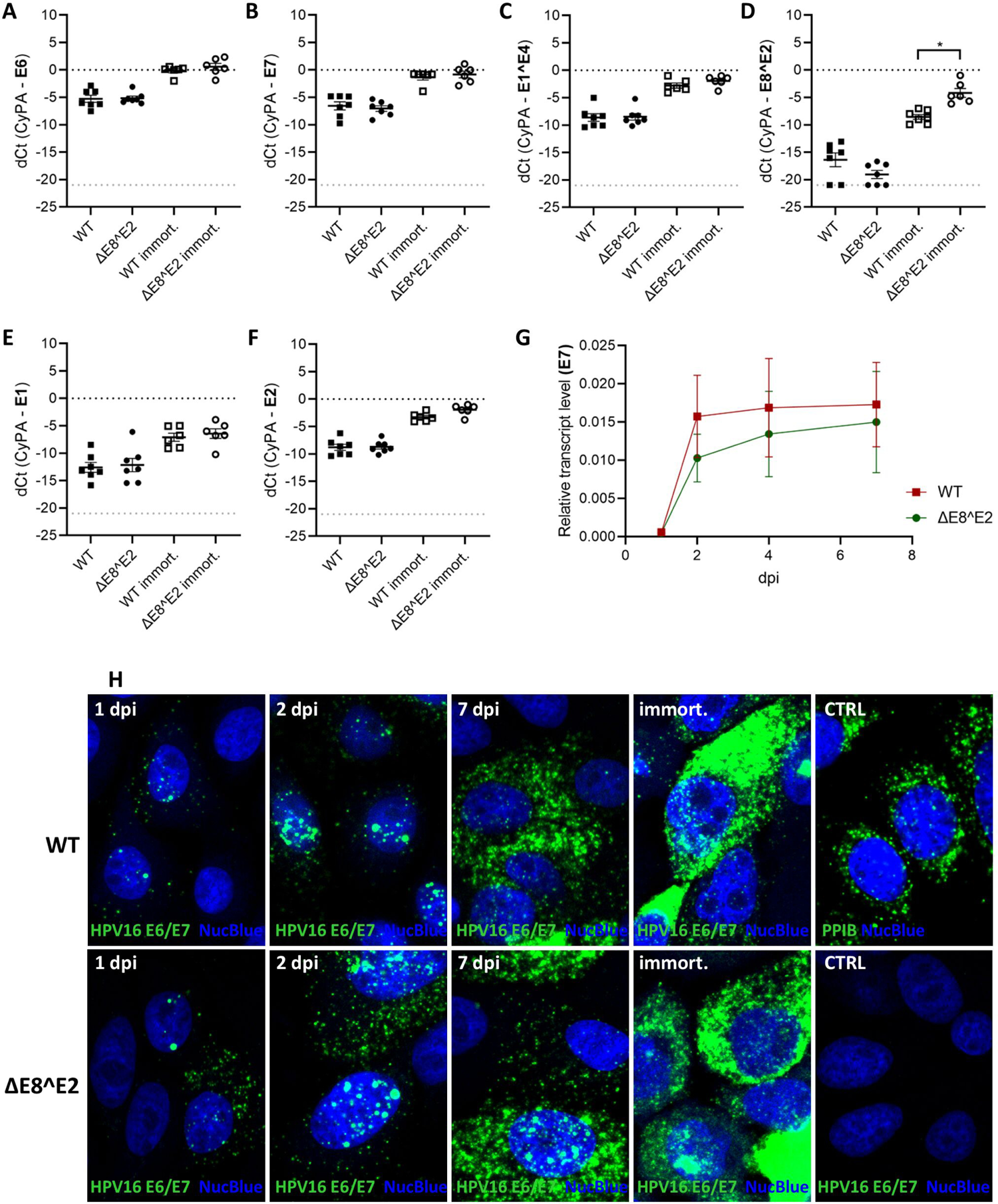
Expression of viral transcripts in HPV16 WT, and ΔE8^E2 mutant infected HFK early after infection. E6, E7, E1^E4, E8^E2, E1, E2 levels at 2 dpi are shown as dCt relative to the housekeeping control gene cyclophilin A (CyPA) (A-F respectively). Viral transcripts in WT-and ΔE8^E2-immortalized cells shown for comparison. Light gray dot line shows limit of detection. Differences statistically significant at *p<0.05. (G) E7 transcript relative to CyPA over 7 dpi. Error bars represent SEM of 7 independent experiments. (H) Representative RNAscope images of infected HFKs growing in monolayer culture at 1, 2, 7 dpi and WT- and ΔE8^E2-immortalized. Uninfected cells shown as positive (PPIB, cyclophilin B) and negative (HPV16 E6/E7) control.

**FIG 7.**
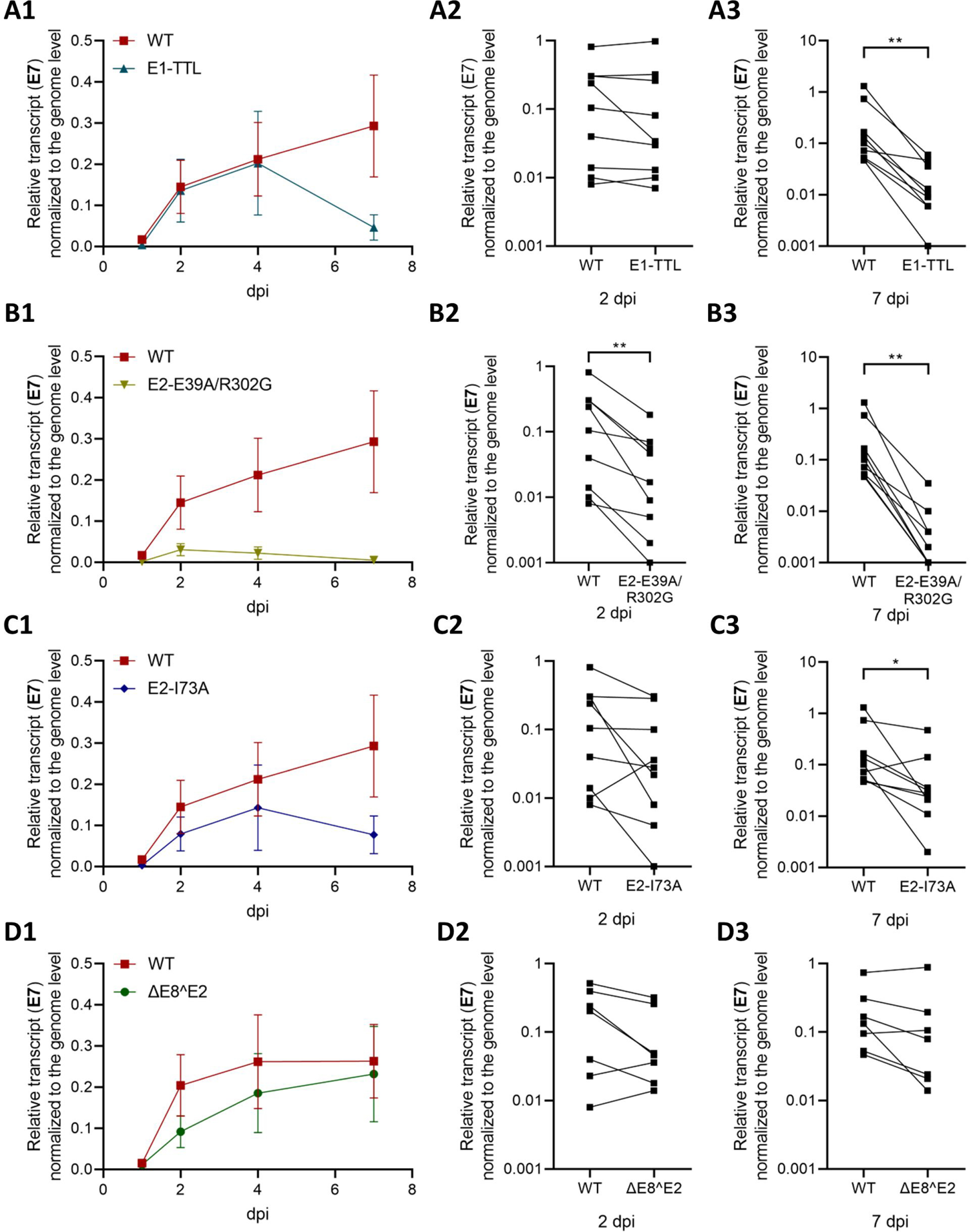
Viral transcripts (E7) normalized to the genome level in HPV16 WT, E1-TTL, E2-E39A/R302G, E2-I73A, and ΔE8^E2 mutant infected HFKs over 7 days in monolayer culture (A1-D1). Error bars represent SEM of 9 (A1-C1) or 7 (D1) independent experiments. Comparison of E7 level normalized to HPV16 DNA between WT and E1-TTL, E2-E39A/R302G, E2-I73A, and ΔE8^E2 mutants at 2 dpi (A2, B2, C2, and D2 respectively) and at 7 dpi (A3, B2, C3, and D3). Differences statistically significant at *p<0.05 and **p<0.01.

### Involvement of E2 protein in regulation of early HPV16 transcription

To further explore the role of E2 in transcriptional regulation immediately after infectious entry, we normalized viral transcript levels to genome level over time post infection. At 2 dpi, early transcript levels normalized to genome copy number was similar in WT and E1-TTL infected HFK suggesting that E1 has no role in transcriptional regulation, as expected (Fig. 7A1 and 7A2). Transcripts were essentially undetectable at 7 dpi with E1-TTL mutant virus due to loss of viral genome (Fig. 7A3). In contrast, p97-derived genome-normalized early transcripts were consistently less abundant in HFK 2 dpi with E2-E39A/R302G when compared to WT (Fig. 7B1, B2). At 7 dpi, most viral genomes and consequently viral transcripts were lost in E2-E39A/R302G infected cells (Fig. 7B3). The results were more complex in E2-I73A infected cells. Even though early transcripts were consistently less abundant compared to WT, it did not reach significance at 2 dpi due to high variability (Fig. 7C1 and 7C2). We again observed high variability at 7 dpi likely due to differences in the level of viral genome loss over time. However, genome-normalized transcript levels were significantly reduced compared to WT (Fig. 7C3).

### HPV16 genome maintenance during long-term monolayer culture of infected keratinocytes

Primary foreskin keratinocytes infected with HPV16 WT, E1-TTL, E2-E39A/R302G, E2-I73A, and ΔE8^E2 as well as uninfected controls were subjected to long-term culture to monitor for viral genome, transcript and cell survival. The infected cells were cultured in the presence of the Rho kinase inhibitor Y27632 up to 7 dpi, after which media without inhibitor was used. Cells were kept in culture until they were still dividing/alive or viral genome and transcript level reached plateau. Viral genome (E6) and transcripts level (E7) was assessed using qPCR. In keratinocytes infected with WT and ΔE8^E2 viruses, we observed a slow increase in viral DNA and transcripts level for up to 5 weeks after which stable viral DNA and transcript levels was detected (Fig. 8A, C, E, G). During this time, both, WT and ΔE8^E2 infected cells immortalized. Viral DNA and transcripts in cells infected with E1-TTL and E2-E39A/R302G decreased dramatically over time and cells senesced approximately 3 weeks after infection (Fig 9A, B). Loss of both viral genome and transcripts was slower in keratinocytes infected with E2-I73A mutant viruses (Fig 9A, B). Loss of HPV16 genomes because of lack of functional E1 and E2 proteins prevented immortalization of infected keratinocytes. In some cases, integration of viral genome was observed in E1 and E2 mutant infected cells resulting in constant detection of viral DNA and mRNA over time. These events were confirmed by exonuclease V-qPCR-based assay (34, 35) and experiments where viral DNA (E6) resistance to exonuclease V digestion was lower than 20% were excluded from further analysis. Viral DNA level in cells infected with ΔE8^E2 was ∼7 times higher than WT at 5 weeks post infection and this difference increased up to 20 times at 8 weeks (Fig 8A). This confirms a role of E8^E2 protein in regulation of viral genome amplification during maintenance phase in undifferentiated keratinocytes. Viral DNA integration was observed in both cases over time; however, ΔE8^E2 integrated into host genome much faster than WT, probably due to presence of excessive number of viral genomes (Fig 8B).

**Fig 8.**
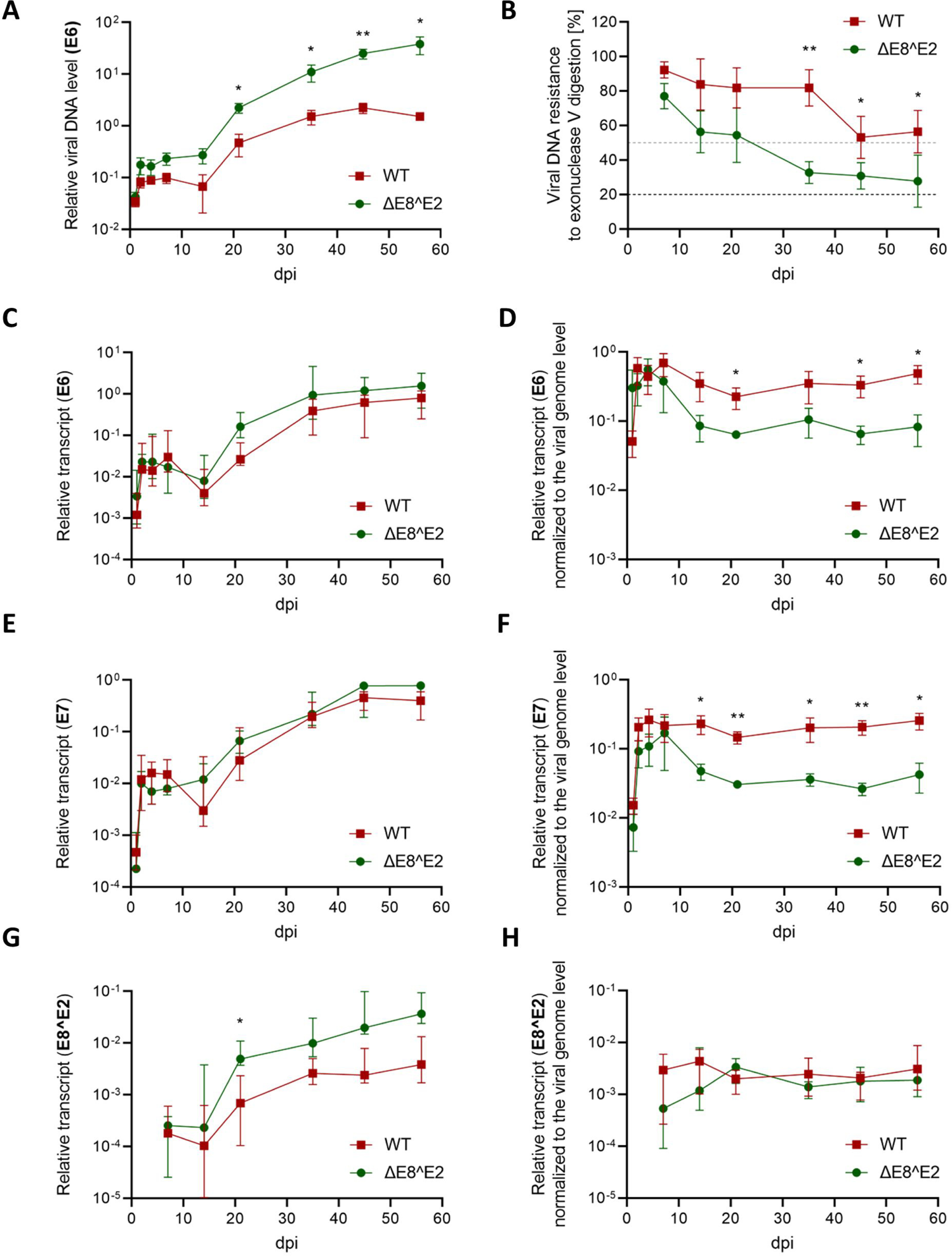
(A) Relative viral DNA level in cells infected with HPV16 WT and ΔE8^E2. (B) Viral DNA resistance to exonuclease V digestion as indicator of episomal/integrated status of genomes. Values below 20% (black dot line) show fully integrated genomes. Values above 50% (gray dot line) show mostly episomal genomes. Viral E6 (C) E7 (E), and E8^E2 (G) transcript in cells infected with HPV16 WT and ΔE8^E2. Transcripts normalized to the viral genome level (D, F, and H). Error bars on graphs C-H represents SEM of 6 experiments. Differences statistically significant at *p<0.05 and **p<0.01.

**FIG 9.**
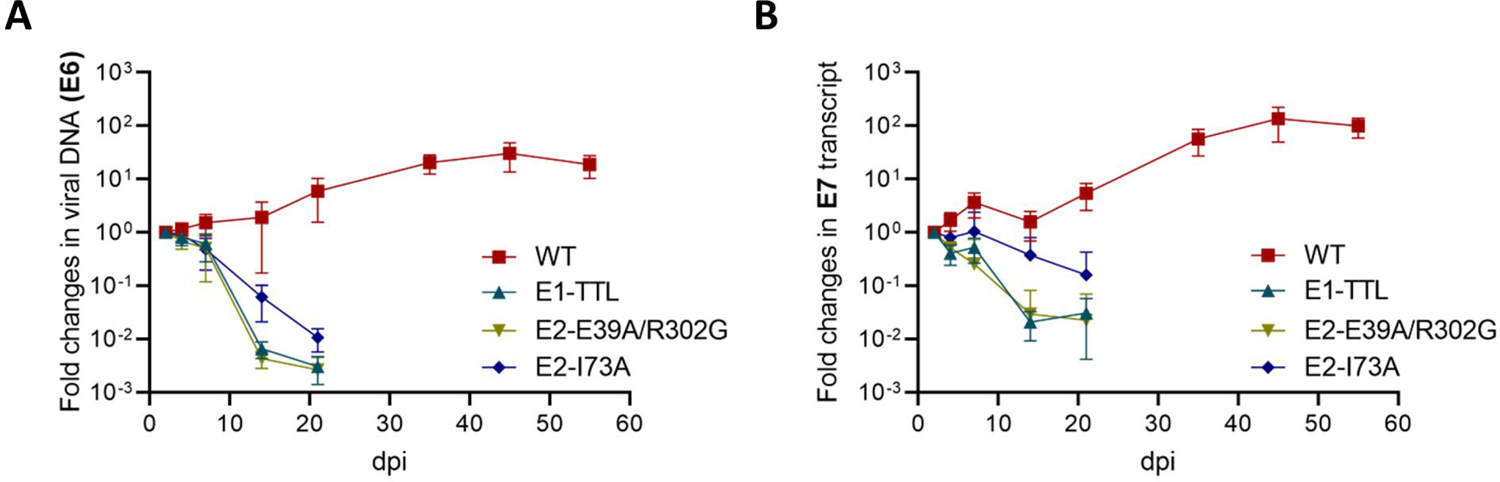
Fold changes in viral genome (E6) (A) and transcript level (E7) (B) during long-term monolayer culture of HFK infected with HPV16 WT, E1-TTL, E2-E39A/R302G and E2-I73A mutants. The data shown on graphs are fold changes relative to results for 2 dpi, normalized to cyclophilin A. Error bars represent SEM of 5 experiments.

### Regulation of viral gene transcription in long-term culture of infected cells

To access the role of E8^E2 in regulation of viral early and pE8 promoters during maintenance in undifferentiated keratinocytes, we analyzed p97 (E7) and pE8 (E8^E2) transcript levels over 8 weeks post infection of HFK with WT and ΔE8^E2 mutant virus. mRNA level was assessed using RT-qPCR and normalized to the cellular gene cyclophilin A. We did not observe significant differences in early HPV16 transcript (E6, E7) (Fig 8C, E). When normalized to viral genome copy numbers, p97 derived early transcript levels were higher in WT infected cells. pE8 promoter derived transcripts were first detected at 2 dpi but values were very close to our qPCR limit of detection (Fig 6D). At later time points, we observed lower level of E8^E2 transcripts in WT infected cells compared to ΔE8^E2 (Fig 8G). However, when normalized to genome copy number, no significant differences were observed (Fig 8H).

### Genome copy number regulation during late stages of the HPV16 lifecycle

Regulation of viral genome amplification under differentiation conditions was evaluated in organotypic raft cultures. Primary human foreskin keratinocytes were infected with HPV16 WT, E1-TTL, E2-E39A/R302G, E2-I73A, and ΔE8^E2 mutant, and cultured for 5-7 dpi in monolayer culture. Next, they were seeded on collagen gel surfaces, and lifted to an air liquid interface on grids. At 21 days post rafting, rafts were collected for *in situ* DNA hybridization, as well as DNA and RNA isolation. Viral genome and transcript level were assessed by qPCR (Fig. 10A) and RT-qPCR (Fig. 10B-D), respectively, and correlated to values obtained from monolayer culture. We observed an increase of viral DNA and mRNA in rafts originated from WT and ΔE8^E2 infected keratinocytes and a decrease in rafts derived from E1-TTL and both E2 mutants compared to monolayer cultures (Fig 10E). HPV16 genomes were undetectable in E1-TTL and E2-E39A/R302G mutant infected cells using DNA scope methodology, because of loss of viral DNA early after infection (Fig 10E). HPV16 E2-I73A genomes were gradually lost during cell division in monolayer culture and in raft tissue, eventually they were undetected by qPCR and DNA scope (Fig 10A, 10E). Genome copy numbers in ΔE8^E2 infected raft tissues were 3.5 times higher than in WT containing rafts. HPV16-specific DNA scope showed much stronger signal in ΔE8^E2 infected 3D cultures confirming an increased level of viral genomes in the absence of functional E8^E2 (Fig 10E). These findings again suggest that E8^E2 protein negatively regulates differentiation induced HPV16 genome amplification. Viral transcripts were increased in raft tissues derived from HPV16 WT and ΔE8^E2 compared to monolayer cell culture but were significantly reduced in tissues derived from keratinocytes infected with E1-TTL and E2 mutants (Fig 10B-D). As expected, viral late transcript E1^E4 and L1 were higher in ΔE8^E2 than WT rafts, however early transcript (E7) was similar in both tissues (Fig 10B-D).

**FIG 10.**
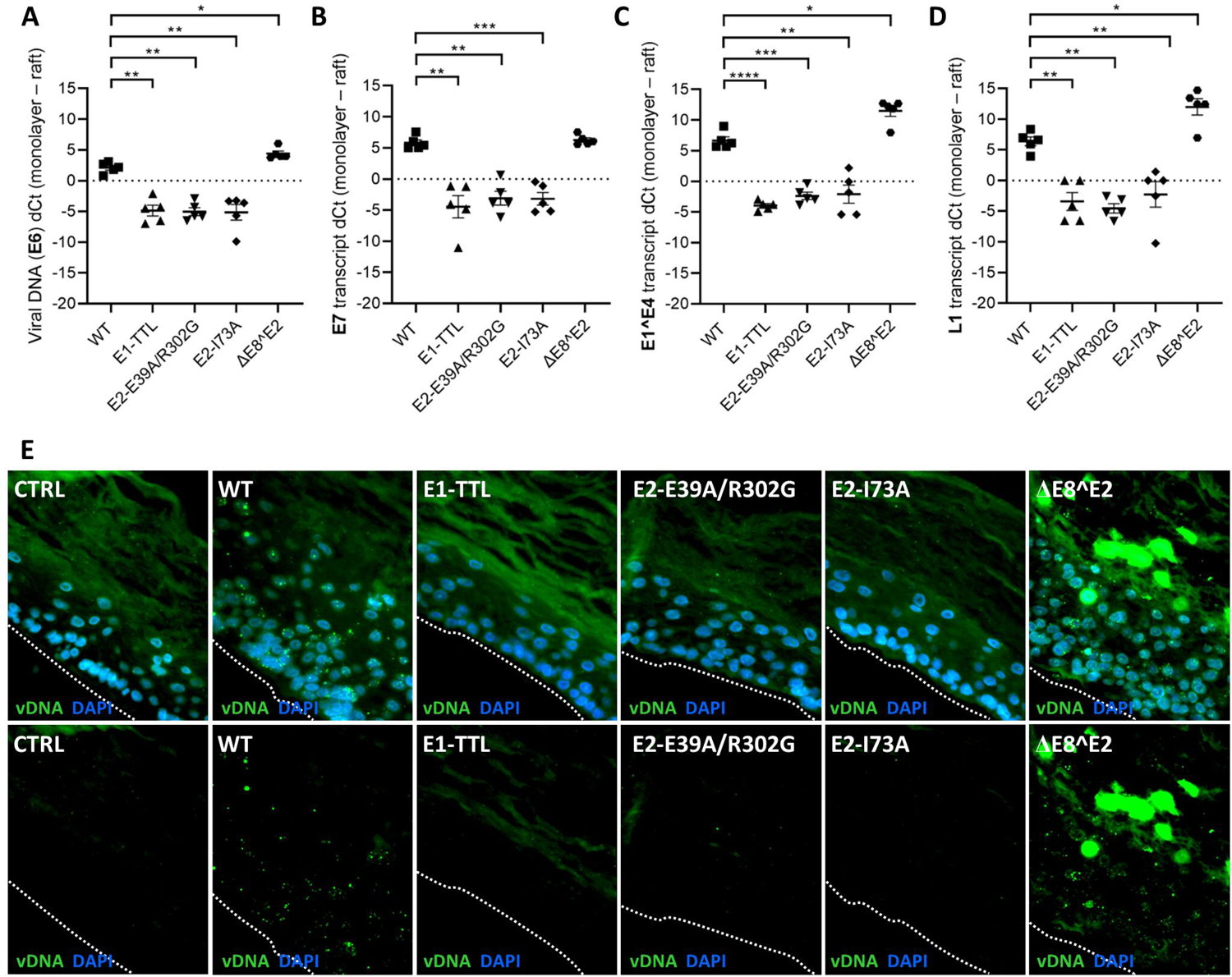
HPV16 genome and transcripts in organotypic raft cultures of keratinocytes infected with HPV-16 WT, E1-TTL, E2-E39A/R302G, E2-I73A, and ΔE8^E2 mutants. (A) Viral DNA (E6), and E7, E1^E4, and L1 transcripts (B-D respectively) expressed as dCt between monolayer at 5-7 dpi and raft culture. Error bars represent SEM of 5 rafts. Differences statistically significant at *p<0.05; **p<0.01; ***p<0.001; ****p>0.0001. (E) Detection of HPV16 DNA by highly sensitive *in situ* hybridization. White dot line underline raft basal layer.

## DISCUSSION

In the present study, we demonstrated for the first time immediate amplification of HPV16 genome after infectious delivery (Fig 1, 3, and 4). It has been generally accepted that viral genome must undergo several rounds of DNA replication after reaching the nucleus in order to establish infection, and genome copy number remains constant during maintenance (4). However, this event has never been experimentally supported due to the lack of an efficient infection model. Most research focused on transient replication of transfected established cell lines or primary keratinocytes cultures in monolayer or 3D cultures (6, 15, 26–32). These systems provided valuable information but were not suitable to study the initial events after infectious delivery of viral genome to the nucleus of primary keratinocytes. Our recently described highly effective infection model, ECM-to cell transfer of HPV16 quasiviruses to primary human foreskin keratinocytes (33), allowed us to track immediate events after infection, such as viral DNA amplification, transcription from early promoter and regulation of these processes without the necessity of selection for immortalized cells. Since the infection model is amenable to a genetic viral screen, we were able to explore the role of transcription and replication functions of E1, E2 and E8^E2 during establishment of infection and for genome copy number control.

We confirmed that viral DNA is amplified after infectious delivery by labeling nascent DNA with the thymidine analogue, 5-ethynyl-2’-deoxyuridine (EdU) (Fig 3C, D) and by immunofluorescent detection of replication foci using DNA scope (Fig 1C, 4C). Viral genome amplification was dependent on the presence of E1 and E2 proteins retaining their replication functions. While EdU labeling only supported the conclusion that viral genome is replicated immediately after nuclear delivery, the observed increase in size of replication foci via DNA scope confirmed amplification. Using EdU labeling, Reinson et al. recently investigated HPV18 DNA replication over cell cycle phases (30) and observed incorporation of EdU into genomes in S and G2 phase three days post transfection of U2OS osteosarcoma cells. Viral replication was restricted to S phase only at later times (30). While informative, these study conditions do not mimic events occurring during natural infection and did not investigate directly viral genome amplification. Our DNA scope analyses revealed the presence of viral genome in the cytoplasm of infected cells, which became more prominent at later times post infection. We were able to exclude that the cytoplasmic signal is due to unspecific staining or detection of viral mRNAs. We demonstrated in the accompanying paper (Bienkowska-Haba et al.) that HPV DNA is amplified during S and G2 phase and genomes which are not associated with chromosomes are lost to the cytoplasm during mitosis. Cytoplasmic DNA is subsequently removed by lysosomal degradation in G1 and S phase and nuclear DNA undergoes the next round of unlicensed amplification. We concluded that this cell cycle dependent cycling of amplification – loss – degradation is responsible for viral genome copy number during maintenance stage.

Our data indicate that replication is not required for early promoter driven transcription. Furthermore, E1 protein does not seem to be involved in p97 promoter regulation as we see similar transcript levels within the first two days post infection with E1-TTL and WT virus (Fig 5A-B, 7A). However, our data support the notion that the transactivation function of E2 protein is essential for efficient transcription from the early promoter. Full-length E2 has been shown to both, activate or repress papillomavirus gene expression based on the abundance and cellular context such as availability and recruitment of cellular factors (8, 36). Mutations in conserved, surface residues of E2 transactivation domain modify transcriptional or replication activity of this protein; however, the phenotype of this mutation may be different in different papillomaviruses (revised in: (8)). The two mutations used in this current study were chosen for separation of replication (E2-E39A) and transcription (E2-I73A) activity of E2 protein (8). Although mutation in E2-E39 position was associated with a decrease in viral replication in HPV16 (13, 37), this function was preserved in BPV (8), HPV31 (38) and HPV11 (12). Most of reporter assays showed that amino acid change of E2-E39 stimulated transcription to the same level as wild-type E2 (13, 36, 38). Our HPV16 E2-E39A/R302G mutant was defective in its replication function (Fig 1). In addition, it was likely unable to bind to DNA and therefore defective in transcriptional transactivation since residue the corresponding R309 of HPV31 has been shown to be important for non-specific interactions with DNA (16). Infection with this mutant virus produces significantly lower amounts of early transcripts at 2 dpi (Fig 7B). A role for E2 in early promoter activation is supported by the analysis of the E2-I73A mutant virus. E2-I73A mutant virus establishes infection; the viral genome is amplified but consistently produced lower levels of early transcripts at 4 and 7 dpi (Fig 7C). Interestingly, E2-I73A mutant genome is slowly lost over time (weeks) resulting in progressively lower viral transcript levels until the cells senesce approximately 3 to 4 weeks post infection. These findings suggest that E2-I73A may be defective for genome tethering and segregation during mitosis even though we do not yet have direct evidence for this notion. Interactions between E2 domain and cellular proteins enabling genome binding to chromosomes was reported in the literature and transactivation domain residue I73 played the role in this process among others, in HPV16, 18 and 31 (8, 11, 37). Involvement of I73 domain in transcription regulation was reported before, and regulation of transcription was seen as mostly independent from interactions of E2 with Brd4 (11, 13, 36, 38).

Our analysis of HFK infected with ΔE8^E2 mutant virus supported the role of E8^E2 in genome copy number regulation, which was previously reported for several papillomaviruses, such as BPV (39, 40), HPV5, 8, 11, 18 (41), HPV31 (20), HPV16 (18, 21, 42). Studies published so far showed that knock out of E8^E2 was associated with 10-100-fold higher genome level compared to wild type. This data comes from short-term assays in transfected normal or immortalized keratinocytes or U2OS osteosarcoma cell line (17). In our time course analysis, we observed a continuous accumulation of ΔE8^E2 DNA over WT (Fig 8A). This suggests a continuous loss of copy number control regulation and confirms that E8^E2 is a negative regulator of HPV16 replication during the maintenance stage of the viral life cycle in undifferentiated keratinocytes. Increased genome copy numbers of E8^E2 genome was also observed under differentiation conditions (Fig 10A, E) consistent with findings of Straub et al. (18). However, whether this is due to higher genome copy numbers at the onset of differentiation or reflects increased differentiation-induced viral genome amplification is unclear and requires further investigation. We consistently observed faster integration of ΔE8^E2 genomes compared to WT HPV16 genomes (Fig 8B) pointing to a higher selection pressure. Similar results has been obtained earlier for HPV31, but not for HPV16 (19, 20), however baseline plasmid amplification of HPV31 was much higher than HPV16 ΔE8^E2 (19).

E8^E2 protein has been described to function as a negative regulator of early promoter activity in undifferentiated cells (19, 20, 43–45) as well as the late promoter under differentiation conditions (18). As E8^E2 shares the DNA-binding and dimerization domain with full length E2 protein, one possible explanation of its mechanism of action is competing with full length E2 for the E2 binding site in viral LCR and blocking or modulation of E2 dependent replication and transcriptional regulation (17, 18). Inhibition of viral replication and transcription was also explained by formation of non-functional E2/E8^E2 heterodimers and/or recruitment of the NCoR/SMRT corepressor complex (19, 22, 43). Additionally, E8^E2 was found to inhibit recruitment of RNA polymerase II and its phosphorylated form to the HPV18 E6/E7 promoter in HeLa cells (46). In our work, we observed similar p97 transcript levels in the presence or absence of the E8^E2 (Fig. 8C, E). However, when normalized to genome copy number, E6 and E7 transcripts were statistically significant less abundant in E8^E2 mutant virus infected cells (Fig. 8D, E respectively). Similarly, E8^E2 transcript levels were increased in E8^E2 mutant virus infected cells thus confirming previous observations by the Stubenrauch group (17, 21). However, no measurable differences were observed when E8^E2 transcript levels were normalized to genome copy number (Fig 8G and H). These data indicate that E8^E2 does not regulate its own promoter by a negative feed-back loop but instead may activate p97 promoter activity. An alternative explanation for our findings is that not all genomes are transcriptionally active. Differences in the transcription profile of WT and ΔE8^E2 may also be a result of different regulation of viral gene transcription from mostly episomal (WT) vs integrated genomes (ΔE8^E2). However, we propose another hypothesis, assuming that E8^E2 protein is involved in early promoter regulation by three-dimensional chromatin organization mediated by YY1 (Ying Yang 1) and CTCF (CCCTC-binding factor) interactions. Both proteins are involved in transcriptional regulation by binding to chromatin and preventing interactions between promoter and nearby enhancer or silencer (47). YY1 is known as a strong repressor of papillomaviruses transcription acting by recruitment of Polycomb repressive complex 1 and 2 (48, 49). Multiple YY1 binding sites in HPV16 genome are located in LCR partially overlapping with E2 binding sites (49, 50). CTCF is a widely expressed DNA-binding protein involved in long-range chromatin interactions and its binding sites located in late region are common for 80% of low and high-risk papillomaviruses. CTCF binds also to HPV16 and 18 sequences located in the E2 open reading frame (51). Mutation of this site caused increased transcription of viral early genes without affecting amplification or maintenance of HPV18 episomes in undifferentiated cells (48, 49, 51). Chromatin loop formation due to the interaction of YY1 and CTCF was shown to repress HPV18 early promoter activity in undifferentiated keratinocytes. (48). We propose that E8^E2 binding to E2 consensus sites located in the LCR may displace YY1 thus disrupting the chromatin loop and resulting in increased p97 promoter activity. We are currently in the process of testing this hypothesis.

## MATERIALS AND METHODS

### Cells

Human embryonic kidney 293TT cells were obtained from John Schiller (NIH, Bethesda, MA) and cultured in DMEM supplemented with 10% FBS, non-essential amino acids, antibiotics and GlutaMAX (Gibco). The same media was used to maintained NIH 3T3 J2 fibroblasts. Human keratinocytes HaCaT cells were purchased from the American Type Culture Collection (ATCC) and grown in low glucose DMEM containing 5% FBS, antibiotics and 2.5 µg/ml of plasmocin (InvivoGen). UMSCC-47 cells (Millipore Sigma, SCC071) were cultured in DMEM with GlutaMAX (Gibco, 1 g/L D-glucose, 110 mg/L sodium pyruvate) supplemented with 10% FBS. UMSCC47 is a head and neck squamous carcinoma cell line isolated from the primary tumor of the lateral tongue of a male patient. It contains 15-18 copies of type II integrated HPV-16 (52, 53). Human neonatal primary foreskin keratinocytes (HFK) were purchased from ATCC (PSC-200-010). We used cells derived from four different donors. HFK were maintained in E media containing 5% FBS and 5 ng/ml of mouse epidermal growth factor (EGF, BD Biosciences; 354010) and 10 µM Rho kinase inhibitor (ROCK) Y-27632 which is known as a factor increasing HFKs lifespan (54, 55). The ROCK inhibitor was excluded 7 days post infection in all experiments involving long-term culture. Immortalized human keratinocytes containing HPV16 episomes were obtain from long term culture of wild type infected HFK. Mitomycin-treated mouse NIH 3T3 J2 fibroblasts were used as feeders for HFKs as described previously (56). Before harvesting the cells for RNA or DNA isolation, feeders were removed by short trypsin treatment and PBS wash.

### Quasivirus generation

Quasivirions were generated using 293TT cells following the improved protocol of Buck and Thompson (57) with minor modifications. The pSheLL16 L1/L2 packaging plasmid was a kind gift from John Schiller (NIH, Bethesda, MA). The plasmid pEGFP-N1 containing the entire floxed HPV16 genome (pEGFP-N1-HPV16) and pBCre plasmid have been described previously (58). 293TT cells were first co-transfected with the pSheLL16 L1/L2 and pEGFP-N1-HPV16 plasmids and 24 hours later transfected with the pBSCre plasmid. After additional two days of culture, cells were harvested and viral particles were purified as described previously (57, 59). Prior to purification, samples were treated with benzonase to remove free DNA. Activity of the Cre recombinase generates two circular plasmids of packable size (pEGFP-N1 and HPV16 genome) therefore isolated viral particles comprise a mixture of pseudovirions, serving as internal non-amplifying control (pEGFP-N1 plasmids) and quasivirions (HPV16 genomes). Viral genome equivalents (vge) were determined by real-time quantitative PCR (qPCR) of encapsidated DNA isolated using the NucleoSpin Blood QuickPure (Macherey-Nagel; 740569.250). To generate mutated quasiviruses, site-directed mutagenesis on the backbone of pEGFP-N1-HPV16 plasmid was used. We created an E1 knock down mutant by introducing a single nucleotide substitution 892G/T into E1 ORF as previously described (24) resulting in the presence of TTL (translation termination linker). Introducing of single nucleotide substitution 2871A/C (13) together with 3659A/G resulted in double mutation in E2 protein, E39A in transactivation domain and R302G in DNA binding domain. The E2-E39A/R302G mutant carries also D435G mutation in E1 protein but this substitution is present outside conservative sequence of protein. E2-I73A plasmid was obtained from Jason M. Bodily, LSUHSC, Shreveport, LA (10). ΔE8^E2 mutant was generated by replacing of single nucleotide (1281T/G) in E8 open reading frame resulting in the presence of stop codon in E8 and silenced mutation in the overlapping E1 gene (18, 19). All plasmids used for quasivirus generation were sequenced in full length.

### Infection using extracellular matrix (ECM)-to-cell transfer

Infection was performed as was published before (33, 60). HaCaT cells were used for secretion of ECM. They were seeded on 60 mm cell culture dishes and grown for 24-48 h until were confluent. Then, they were incubated in PBS supplemented with 0.5 mM EDTA for up to 2 h in order to remove the cells. To prevent outgrowth of any residual HaCaT cells, the dish surface was treated with 8 μg/ml mitomycin for 4 h. Optiprep-purified viral particles (1-2 × 10^8^ vge/dish, around 100-200 vge/cell) diluted in 1.5 ml E medium were added to the ECM for 2-3 h at 37°C. Next, low passage primary keratinocytes were added in the number appropriate for each time point. After at least 6 h post seeding of HFKs, 1×10^5^ mitomycin-treated J2 fibroblast feeder cells were added. Infection was continued depending on experiment up to 1-, 2-, 4-, and 7-days post infection. At the time of the collection infected cells reached around 70-80% of confluency. Long-term cultures were carried up for up to 56 dpi and HFKs were split and collected for subsequent DNA and RNA analysis every 7-14 days.

### RNA isolation and RT-qPCR

Total RNA from HFK was extracted using the RNeasy Plus Mini RNA Isolation kit (74236; Qiagen). RNA from raft tissues was extracted using the Qiagen QIAcube via the miRNeasy aqueous-phase protocol. Isolated RNA samples were subsequently treated with DNase I (M0303L; NEB) prior to reverse transcription; between 0.5 to 1 µg of total RNA was reverse transcribed into cDNA using an ImProm II Reverse Transcriptase kit (Promega). Equal amounts of cDNA were quantified by RT-qPCR using the IQ SYBR green Supermix (Bio-Rad) and a CFX96 Fast Real-Time system (Bio-Rad). PCRs were carried out in triplicates, and transcript levels were normalized to cyclophilin A (CyPA). Mock reverse transcribed samples were included as a negative control. Primer sequences used in current work are listed in TABLE 1. Bio-Rad CFX Manager 3.1 and Bio-Rad CFX Maestro 1.1. software was used to analyze the data.

**TABLE 1.**
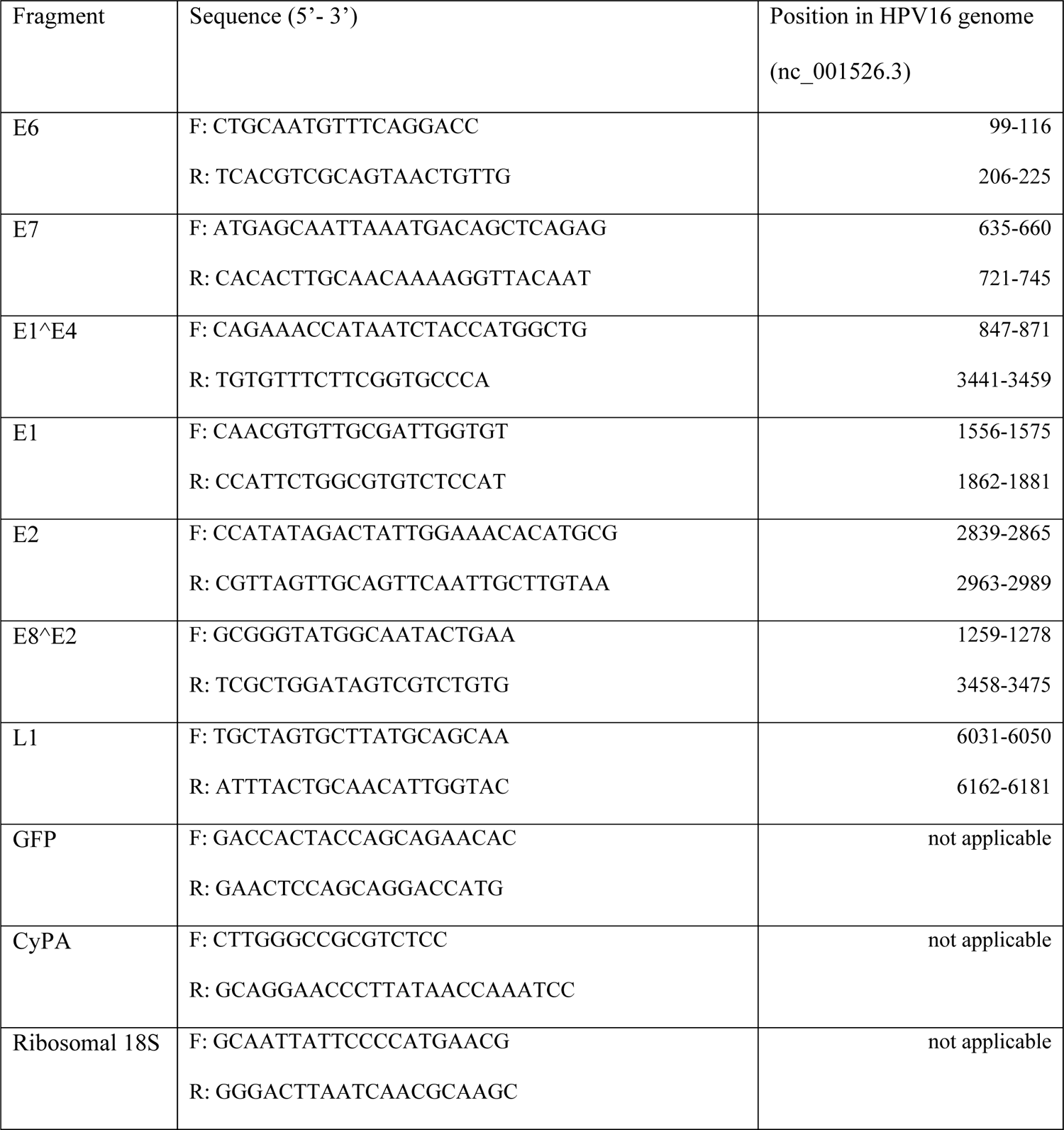
Oligonucleotides used in qPCR

### DNA isolation and qPCR

DNA from monolayer cultures as well as from rafts was isolated using the NucleoSpin Blood QuickPure kit (740569.250; Macherey-Nagel) according to the manufacturer’s instructions with some modifications for raft tissues. Fragments of raft up to 25 mg were placed in 200 µl of PBS, 200 µl of BQ buffer and 30 µl of proteinase K, incubated at 70°C for 1h, then centrifuged for 5 min at 10,000 rpm. Supernatants were used for subsequent steps as recommended by manufacturer. One hundred nanograms of DNA was quantified by RT-qPCR using the IQ SYBR green Supermix (Bio-Rad) and a CFX96 Fast Real-Time system (Bio-Rad). Genome level was assessed based on E6 (primers as above) normalized to cyclophilin A. The Bio-Rad CFX Manager 3.1 and Bio-Rad CFX Maestro 1.1 software were used to analyze the data.

### Detection of nascent viral DNA immediate/early after infection

Shortly after infection of HFKs, newly synthesized DNA was labeled with 5-ethynyl-2’-deoxyuridine (EdU). EdU-DNA pull down was performed using Click-iT® Nascent RNA Capture Kit (C10365, Moleular Probes, Life Technologies), modified to operate with labeled DNA. Primary keratinocytes were infected through ECM-to cell transfer and, 10 µM EdU was added to the media around 12-16 h after infection. HFKs were then cultured in/without the presence of EdU up to 24 or 48 h post infection, then they were collected, and DNA was isolated as shown above. Between 2-5 µg of EdU-DNA or unlabeled DNA was biotinylated with 0.75 mM biotin azide (PEG4 carboxamide-6-azidohexanyl biotin) using Click-iT reaction carried out for 50 min at room temperature. Biotinylated DNA was then purified by ethanol extraction and streptavidin magnetic beads (Dynabeads MyOne Streptavidin T1) were used to capture the newly synthetized pool of DNA. EdU-DNA as well as input DNA was quantified by RT-qPCR using the IQ SYBR green Supermix (Bio-Rad) and a CFX96 Fast Real-Time system (Bio-Rad). Two pairs of primers were used, detecting viral E6 and cellular cyclophilin A (CyPA). Nascent DNA level was expressed as difference between CT values of DNA pulled down from unlabeled and labeled samples (dCt*) normalized to input DNA: dCt* = unlabeled DNA dCt (pull down – input) – EdU labeled DNA dCt (pull down – input).

### Organotypic raft culture

Organotypic raft cultures generated from HFK infected for 5 to 7 days with HPV16 quasivirions were grown as described (56, 61). One million keratinocytes were seeded onto the surface of the collagen gel containing fibroblasts feeders. Following attachment, the gel with keratinocytes layer was lifted and placed onto stainless steel grid in a culture dish. Culture medium was added to the dish so that the keratinocyte/collagen plug was exposed to the air from above and to the medium below. The medium was changed every other day maintaining the air-liquid interface. Rafts were grown for 21 days and samples were collected for *in situ* DNA hybridization as well as total DNA and RNA isolation. Raft tissues were fixed for 30 min in 4% paraformaldehyde at 4°C, washed three times in cold PBS and once in cold 70% ethanol, and stored in fresh cold 70% ethanol at 4°C until processing and paraffin embedding. Rafts generated from uninfected HFKs seeded on ECM were used as control.

### Detection of the episomal or integrated HPV16 genome

Episomal or integrated status of viral genome was evaluated using an exonuclease V-qPCR-based assay as described before (34, 35) with minor modifications. Briefly, 100 ng of DNA was treated with or without 6.6 units of ExoV (RecBCD, NEB) in a 10 µl reaction volume and incubated for 1 h at 37°C. Digestion was followed by heat inactivation at 95°C for 10 min. DNA isolated from UMSCC47 cells served as a control of integrated HPV16 genome. DNA from 15 ng equivalent was quantified by RT-qPCR using the IQ SYBR green Supermix (Bio-Rad) and a CFX96 Fast Real-Time system (Bio-Rad). Two pairs of primers were used, allowed for detection of viral E6 and human rDNA (18S). Relative DNA amounts were calculated based on a standard curve generated using a 10-fold dilution spanning 5 logs that started at 100 ng of undigested UMSCC47. The percent of DNA resistant to exonuclease digestion was calculated relative to undigested DNA. HPV16 status was recognized as episomal when ExoV-resistance was greater than 20% and integrated when resistance was lower than 10%.

### Detection of HPV16 RNA in HPV16-infected HFKs grown in non-differentiating conditions

Viral RNA detection (**RNAscope**) in infected cells was performed by utilizing the Multiplex Fluorescent v2 kit (catalog number: 323100; Advanced Cell Diagnostics [ACD]) and a probe targeting HPV16/18 E6 and E7 (cat. no. 311121; ACD) according to the manufacturer’s protocol. Briefly, infected and control cells cultured on coverslips were fixed with 4% paraformaldehyde (PFA) for 15 min at room temperature at 2 and 7 dpi. After washing with PBS, samples were dehydrated with 50%, 70%, and 100% ethyl alcohol gradient for 5 min each at room temperature. Then, cells were rehydrated with 70% and 50% ethyl alcohol gradient for 2 min each and finally incubated twice with PBS for 10 min. Next, slides were incubated with pretreatment reagents (cat. no. 322381; ACD): hydrogen peroxide and protease III at room temperature for 10 min each. After a PBS wash, samples were incubated with pre-warmed target probes for 2 h at 40°C. In addition to the probe targeting HPV16/18 E6 and E7, the positive control probe for cellular RNA detection (PPIB, cat no. 313901; ACD) provided by the supplier was used (Fig 2). Signal amplification and detection reagents (cat. no. 323110; ACD) were applied sequentially and incubated at 40°C in AMP1, AMP2, AMP3, and HRPC1 reagents for 30, 30, 15, and 15 min, respectively. Before adding each reagent, samples were washed twice with washing buffer (cat. no. 310091; ACD). For fluorescence detection, Alexa Fluor 488 tyramide (cat. no. T20912; Invitrogen) in RNAscope Multiplex TSA buffer (cat. no. 322809; ACD) was added for 30 min at 40°C. After extensive final washing, slides were air-dried and mounted in ProLong Glass Antifade Mountant with NucBlue Stain containing Hoechst 33342 (cat. no. P36985, Invitrogen) for imaging. Confocal images were acquired as single sections with the Leica TCS SP5 spectral confocal microscope using a 63X objective and processed using Leica Application Suite X software.

### Detection of HPV16 DNA in HPV16-infected HFKs grown in non-differentiating conditions

Viral DNA detection (**DNAscope**) was performed by utilizing a Multiplex Fluorescent v2 kit (cat. no. 323100; ACD) and the probe targeting HPV16/18 L1 (cat no. 315601; ACD). We modified the RNAscope procedure as described by Deleage et al. (62) by adding an RNase tissue pretreatment step followed by a short denaturation. Infected and control cells cultured on coverslips were fixed with 4% paraformaldehyde (PFA) for 15 min at room temperature at 2 and 7 dpi. After washing with PBS, samples were dehydrated, rehydrated and washed in PBS as in RNAscope (see above). Next, slides were incubated with pretreatment reagents (cat. no. 322381; ACD): hydrogen peroxide and protease III at room temperature for 10 min each. Slides were rinsed in Milli-Q water and incubated with RNase A (25 µg/ml, cat no. EN0531; Thermo Scientific) for 30 min at 37°C. The RNase tissue pretreatment step was followed by a short denaturation in which the slides were incubated at 60°C with pre-warmed probes for 15 min and then transferred to 40°C for hybridization overnight. Amplification and detection were performed according to the manufacturer’s protocol, using RNAscope 0.5× washing buffer for all washing steps. Positive control probe for cellular RNA detection (PPIB, cat no. 313901; ACD), together with or without RNase A treatment was used for testing DNA detection specificity (Fig 2). Confocal images were acquired with the Leica TCS SP5 spectral confocal microscope using a 63X objective and processed using Leica Application Suite X program. Quantification of viral DNA puncta in cells was done using ImageJ software.

### Detection of HPV16 DNA in organotypic raft cultures by DNAscope

Paraffin-embedded sections of raft tissues were heated at 60°C for 1 h, deparaffinized in xylene for 10 min, followed by dehydration in 100% ethanol for 5 min before air drying. Next, slides were incubated with RNAscope hydrogen peroxide for 10 min at room temperature to block endogenous peroxidases. Heat-induced epitope retrieval was performed by boiling sections in RNAscope Target Retrieval reagent for 15 min, and the sections were immediately washed in Milli-Q water and then dehydrated in 100% ethanol for 5 min before air drying. Tissue sections were then incubated with RNAscope Protease Plus for 30 min at 40°C. Slides were rinsed in Milli-Q water and incubated with RNase A (25 µg/ml, cat no. EN0531; Thermo Scientific) for 30 min at 37°C. The RNase tissue pretreatment step was followed by a short denaturation in which we incubated the slides at 60°C with pre-warmed probes for 15 min and then transferred to 40°C for hybridization overnight. Amplification and detection were performed according to the manufacturer’s protocol, using RNAscope 0.5× wash buffer for all washing steps. After extensive washing slides were air-dried and mounted in Gold Antifade containing 4’,6-diamidino-2-phenylindole (DAPI) (P3693; Life Technologies) for imaging. Images were captured using a Leica CTR6000 fluorescence microscope with 40X lens and processed with Leica Application Suite X software.

## ACKNOWLEGDEMENTS

We thank Dr. Jason Bodily for providing reagents and helpful discussions. This work was supported by the NIH through grants to M.S. (R01CA211576 and P30GM110703) and R.S.S. (R01DE025565) and by the Feist Weiller Cancer Center.

